# Discovery of PilU as a second Type IV pilus retraction motor in *Myxococcus xanthus*

**DOI:** 10.64898/2026.05.11.724270

**Authors:** Mahdia Rahman, Kalpana Subedi, Andrea Harms, Daniel Wall, Anke Treuner-Lange

## Abstract

Type IV pili (T4P) drive social (S) motility in *Myxococcus xanthus* through cycles of extension and retraction powered by the ATPases PilB and PilT. Although the canonical retraction ATPase PilT is essential for force generation, *M. xanthus* encodes four PilT-like paralogs whose contributions to motility remain unclear. Here, we identify MXAN_1995 as the long-sought PilU protein that serves as a second T4P retraction motor. A frameshift mutation or deletion of *pilU* abolishes S-motility, while preserving pilus assembly and exopolysaccharide (EPS) production, phenocoping the *pilT* mutant. Single-cell analyses revealed that Δ*pilU* mutants exhibit rare, low-force movements, consistent with a role for PilU in force generation. Fluorescence microscopy showed PilU localizes predominantly to cell poles, similar to PilT; this localization is independent of PilT but partially dependent on core T4P assembly proteins. Notably, calcium differentially modulates motility, enhancing movement in wild-type cells while suppressing it in Δ*pilU* mutants, indicating a role for PilU under varying environmental conditions. Structural modeling, together with an intragenic suppressor, highlights a regulatory function for the intrinsically disordered C-terminal region of PilU. Together, our findings establish PilU as a secondary retraction ATPase and uncover a dual-motor retraction system that is environmentally responsive and mechanically tunable.

**Importance:** T4P are widespread motility and adhesion systems that enable bacteria to move, interact, and form multicellular communities. While the primary retraction ATPase PilT is well characterized, the function of additional PilT-like proteins remains unclear in many species. This work provides the first mechanistic characterization of PilU in *Myxococcus xanthus*, a model for multicellular behavior and T4P biology. We show that PilU is essential for productive T4P retraction, functioning as an accessory motor that enhances or stabilizes PilT-driven force generation. We further reveal that PilU activity is modulated by environmental calcium and depends on a flexible C-terminal region that influences motor complex dynamics. These findings uncover a dual-motor architecture that enables adaptive control of T4P retraction in response to environmental cues.

## Introduction

Motility is a key determinant of bacterial fitness, contributing to habitat colonization, virulence, and biofilm formation [1]. One widespread mechanism of bacterial motility is mediated by type IV pili (T4P), dynamic cell-surface filaments that also participate in pathogenicity, DNA uptake, protein secretion, and biofilm development. T4P-driven movement results from repeated cycles of pilus extension, surface attachment, and retraction, generating retraction forces exceeding 100 pN and making T4P among the most powerful biological nanomachines described [2–4].

*Myxococcus xanthus* is a Gram-negative, soil-dwelling bacterium renowned for complex multicellular behaviors such as predation, development, and sporulation, all of which rely on coordinated motility [5]. This organism employs two genetically and mechanistically distinct motility systems. Adventurous (A) motility supports single-cell movement on firm surfaces and depends on Agl/Glt focal adhesion complexes spanning the cell envelope [6]. In contrast, social (S) motility predominates on moist surfaces and is powered by T4P. Secreted polysaccharide referred to exopolysaccharides (EPS) is important for S-motility and can affect T4P extension or retraction [7].

The extension and retraction of T4P are driven by the conserved T4 pilus machinery (T4PM) [8]. In Gram-negative bacteria, this complex spans the outer membrane, periplasm, inner membrane, and cytoplasm and comprises approximately 15 conserved components [9, 10]. The machine consists of a basal body embedded in the cell envelope and an external pilus fiber composed primarily of the major pilin PilA. Basal body subcomplexes include the outer membrane pore, alignment complex, and motor complex. In addition, a priming complex containing PilY1 and several minor pilins facilitates pilus assembly and mediates surface attachment and retraction signaling [9].

T4P dynamics are powered by the motor complex, which includes the platform protein PilC and the ATPases PilB and PilT. PilB and PilT associate with the cytoplasmic base of the T4PM in a mutually exclusive fashion to power T4P extension and retraction, respectively [10]. PilB and PilT belong to the ASCE family of AAA⁺ ATPases and assemble as hexameric rings surrounding a central pore [8, 11–14]. Through interactions with PilC and PilM, ATP binding and hydrolysis induce coordinated conformational changes within the hexamer that are thought to drive PilC rotation and pilus polymerization or depolymerization [15–17].

The rod-shaped *M. xanthus* cells move across surfaces in the direction of their long axis exhibiting distinct leading and lagging cell poles. While the T4PM core is present at both poles, T4P only assemble at one pole at a time, allowing unidirectional movement with a piliated leading and a non-piliated lagging cell pole [5, 18]. Consistent with the unipolar T4P formation, the PilB extension ATPase localizes to the leading cell pole, while PilT predominantly localizes to the lagging pole and occasionally localizes to the leading pole stimulating retractions [19].

Dedicated retraction ATPases are a defining feature of T4P systems. PilT functions as the primary depolymerization motor and is required for high-force, high-velocity pilus retraction [3, 4]. Loss of *pilT* in many bacteria abolishes twitching motility, DNA uptake, and surface-associated behaviors, frequently resulting in hyperpiliation due to continued extension in the absence of retraction [3, 4, 8, 20–22]. In several Gram-negative bacteria, including *Vibrio cholerae*, *Pseudomonas aeruginosa*, and *Neisseria* spp., a second PilT-like ATPase, PilU, is encoded immediately downstream of *pilT* within the same operon [23–25]. PilU shares substantial sequence homology with PilT and exhibits ATPase activity *in vitro*, yet its contribution to motility differs across systems. In some organisms, PilU functions as an accessory or conditional retraction motor, enhancing retraction efficiency under high-load conditions or contributing to sustained force generation [24, 26, 27]. In others, PilU activity is PilT dependent, suggesting a model in which PilU modulates retraction dynamics rather than acting as an independent motor [23, 24]. These observations support a division of labor among retraction ATPases, with PilT providing rapid, high-force retraction and PilU fine-tuning retraction efficiency, processivity, or coordination with environmental cues.

Notably, *M. xanthus* encodes an expanded set of PilT-like AAA⁺ ATPases, including four paralogs in addition to the canonical PilT, with sequence identities ranging from 58% to 66%. This expansion suggests increased regulatory complexity and potential specialization of retraction motors in a bacterium that relies heavily on collective motility. T4P retraction normally generates sufficient force to pull cells forward, with wild-type cells producing ∼150 pN of stall force, yet *pilT* mutants can still generate high velocity retractions with lower force, ∼70 pN, indicating the existence of an additional retraction motor in *M. xanthus* [3, 28]. Here, we identify MXAN_1995 as the long-sought PilU retraction ATPase involved in T4P-dependent social motility. A frameshift mutation or deletion of MXAN_1995 abolishes swarming motility without affecting pilus assembly or EPS production, thus phenocopying a *pilT* mutant. Fluorescence microscopy further reveals that the protein localizes to the cell poles and colocalizes with PilT. Together, these findings establish *MXAN_1995* as the *M. xanthus pilU* gene and support a model in which multiple retraction ATPases cooperate to modulate force generation and pilus dynamics during S-motility.

## Results

### Identification of *MXAN_1995* as the *pilU* gene required for S-motility

To comprehensively identify genetic factors involved in T4P function in *M. xanthus*, we have characterized an extensive collection of unbiased S-motility mutants generated by chemical and UV mutagenesis in strains that lacked A-motility. Each of these 60 mutants were quantified for pili production by transmission electron microscopy [14]. By using this approach, the *pilT* and *dsp*/*dif* genes were identified where their mutations resulted in S-motility defects, although pili were still assembled. Additionally, other mutations were mapped outside of the *pilT* locus, or the mutants did not have a *dsp*/*dif* none clumping phenotype, although pili were still expressed. Our interest focused on this class mutants because they have an implied role in pili function as opposed to pili biogenesis. Through genomic sequencing and validation approaches, two new S-motility genes were identified, *sglS* and *sglT*, and our analysis indicated they play redundant essential roles in T4P function at high temperatures, though they were not homologous to known *pil* genes [29–32]. Here we investigate an additional S-motility mutation in strain DK2120, which contained the *cglB2* A-motility mutation. This S-motility mutation mapped outside of known loci and exhibited a *pilT*-like phenotype, producing pili and forming cell clumps, but also displaying a temperature-sensitive (Ts^−^) S-motility phenotype (Wu et al., 1997).

To identify the causative mutation in DK2120, its genome was sequenced. A single cytosine insertion mutation within a homopolymeric tract of five cytosines near the 3′ end of *MXAN_1995* caught our interest because this ORF was a *pilT* paralog (Fig. 1A). To test our hypothesis that the *MXAN_1995* frameshift mutation caused the S-motility defect, the wild-type (WT) gene was cloned in pMR3487 and integrated into the DK2120 genome. Complementation tests demonstrated that the WT *MXAN_1995* allele restored S-motility (Fig. 1B), confirming that the *MXAN_1995* mutation was responsible for the S-motility defect and, hereafter, was called *pilU*. Comparative genomics found that *pilU* was typically located within a conserved four-gene cluster in the Myxococcales phylum (Fig. 1C), and in *Archangium* and *Melittangium*, it is found in proximity to *pilY1* and genes encoding minor pilins (Fig. S1). Sequence analysis showed that PilU contains the conserved Walker A and Walker B motifs, as well as the Asp and His Boxes characteristic of P-loop NTPases involved in ATP binding and hydrolysis [8, 33]. In addition, PilU possesses a C-terminal ∼112–amino acid intrinsically disordered region (IDR) (Fig. 1A) that is conserved only in myxobacteria.

**Figure 1.**
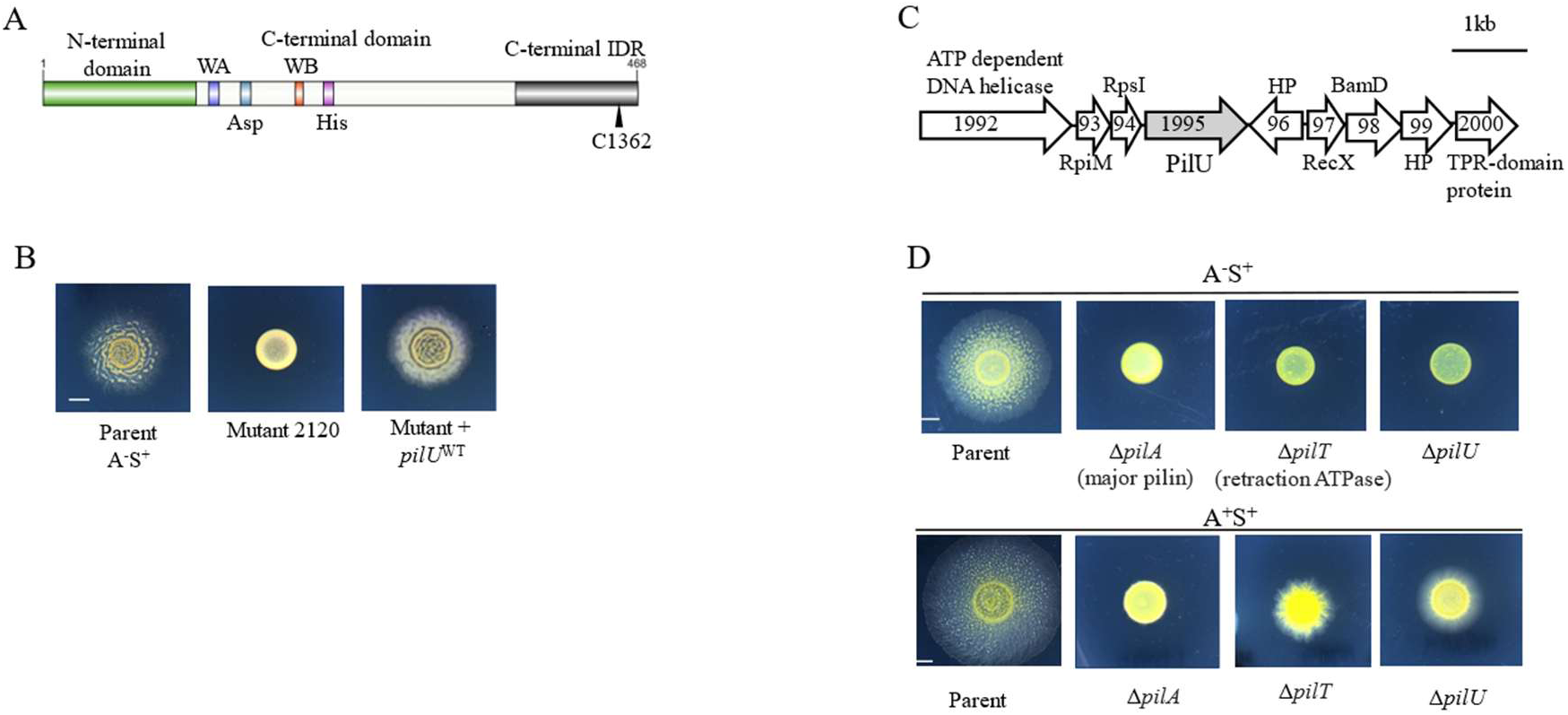
Complementation and genetic context of *pilU*. (A) Domain organization of PilU. Vertical bars indicate four conserved motifs in the ATPase domains: Walker A (WA) box, Asp box, Walker B box (WB), and His box. Black triangle marks the frame-shift mutation in DK2120. (B) Complementation of the DK2120 *pilU-fs* mutant with WT *pilU*. Parent strain DK1218. (C) Genomic organization of the *pilU* locus in *M. xanthus.* Numbers represent MXAN locus tags. (D) Phenotypic analysis of *pilU* deletion mutants in A⁻S⁺ (Parent strain DK1217) and A⁺S⁺ (Parent strain DK1622) backgrounds with Δ*pilA* and Δ*pilT* controls. All S-motility assays done on soft agar with 2 mM CaCl_2_; scale bars, 2.5 mm.

To independently validate the functional role of *pilU* and its null phenotype, we generated an in-frame deletion of this gene in an A⁻S⁺ background (DK1217; *aglQ1*). The resulting Δ*pilU* mutant showed a nonmotile phenotype at 33 °C that was indistinguishable from Δ*pilA* and Δ*pilT* mutants (Fig. 1D, upper panel), consistent with a loss of T4P-dependent motility. However, this Δ*pilU* mutant exhibited a Ts^−^ motility phenotype similar to DK2120 (*cglB2 pilU-fs*), both showing residual motility at lower temperatures (18 °C) (Fig. S2). Deletion of *pilU* in an otherwise WT A⁺S⁺ background (DK1622) resulted in a severe S-motility defect at 33 °C, characterized by limited flares of cells emerging from the colony and markedly reduced spreading on soft (0.5%) agar (Fig. 1D, lower panel). The residual colony expansion, which contrasted with a Δ*pilA* mutation, suggested that the Δ*pilU* mutation, in the presence of A-motility, retains limited soft agar swarming. This phenotype was similar to the Δ*pilT* mutant (Fig. 1D), where we attribute soft agar swarming dependent on A-motility when cells can bundle together from pili/EPS production, hence facilitating soft agar swarming [14, 34]. Together, these results demonstrate that *pilU* was an important component for S-motility and likely functions with the T4P retraction machinery in *M. xanthus*.

### PilU encodes a T4P retraction ATPase

Among the four PilT paralogs in *M. xanthus*, *MXAN_6705* and *MXAN_6706* were highly conserved across Myxococcales, whereas PilU was less conserved (Fig. 2A). Phylogenetic analysis found that PilT and PilU cluster closely together, indicating a close evolutionary relationship (Fig. 2B). Despite differences in sequence conservation, all five paralogs contain the hallmark Walker A and Walker B motifs, as well as the conserved Asp and His boxes characteristic of functional P-loop NTPases (Fig. 2C, S3).

**Figure 2.**
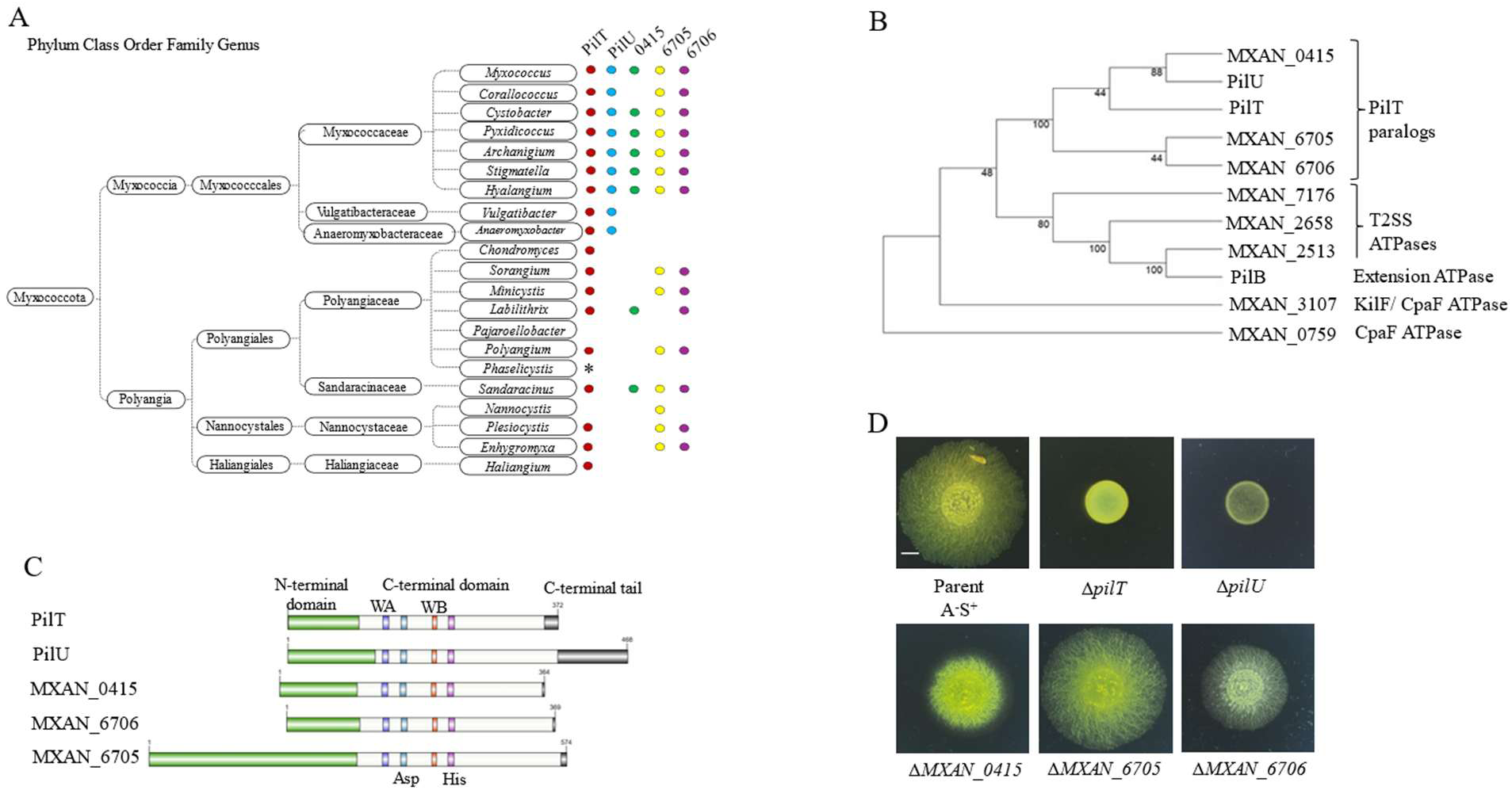
Distribution and phylogeny of PilT paralogs in Myxococcota. (A) Occurrence of *pilT* (*MXAN_5787*), *pilU* (*MXAN_1995*) and three additional *pilT* paralogs across phylum Myxococcota. The presence of paralogs indicated by colored circles (E-value < 10^-10^). Blank, not present; asterisks, unavailable genomes. (B) Phylogenetic tree showing the relationships among hexameric ATPase family in *M. xanthus*. (C) Domain organization of PilT paralogs. The N- and C-termini are conserved except *MXAN_6705* and *pilU* have longer N- and C-termini, respectively, that are intrinsically disordered. Vertical bars indicate four conserved sequence motifs in the ATPase domains: Walker A (WA) box, Asp box, Walker B box (WB), and His box. See Figure S1 for sequence alignment. (D) Swarm phenotypes on soft agar 2 mM CaCl_2_ of indicated strains in A⁻ backgrounds; scale bar, 2.5 mm. Parent strain DK1622 Δ*cglC*.

To investigate the roles of these PilT-like ATPases in S-motility, we generated in-frame deletion mutants in each of the three additional paralogs in a A^ꟷ^S^+^ strain. Unlike the Δ*pilU* mutant, these paralogs produced varying degrees of S-motility defects, with the Δ*MXAN_041*5 and Δ*MXAN_6706* eliciting the strongest phenotypes (Fig. 2D). Based on its phylogenetic relationship to PilT and its critical role in motility, further supports that *MXAN_1995* encodes a *bona fide pilU* gene.

### PilU is dispensable for pilus assembly and EPS production

Since T4P serves as the molecular engine to drive S-motility, we tested whether the Δ*pilU* mutant produced pili. For this, we conducted shear assays where surface exposed thin pili were broken off by shear forces followed by fractionation and immunoblotting with α-PilA antibody. The PilU mutant exhibited about 2.5-fold higher surface exposed PilA comparable to parent strain, indicating that pilus assembly remained intact (Fig. 3A). Similar results were observed with the *pilT* and the *pilT-U* double mutant (Fig. 3A). A *pilA* mutant served as negative control where no sheared pili was detected.

**Figure 3.**
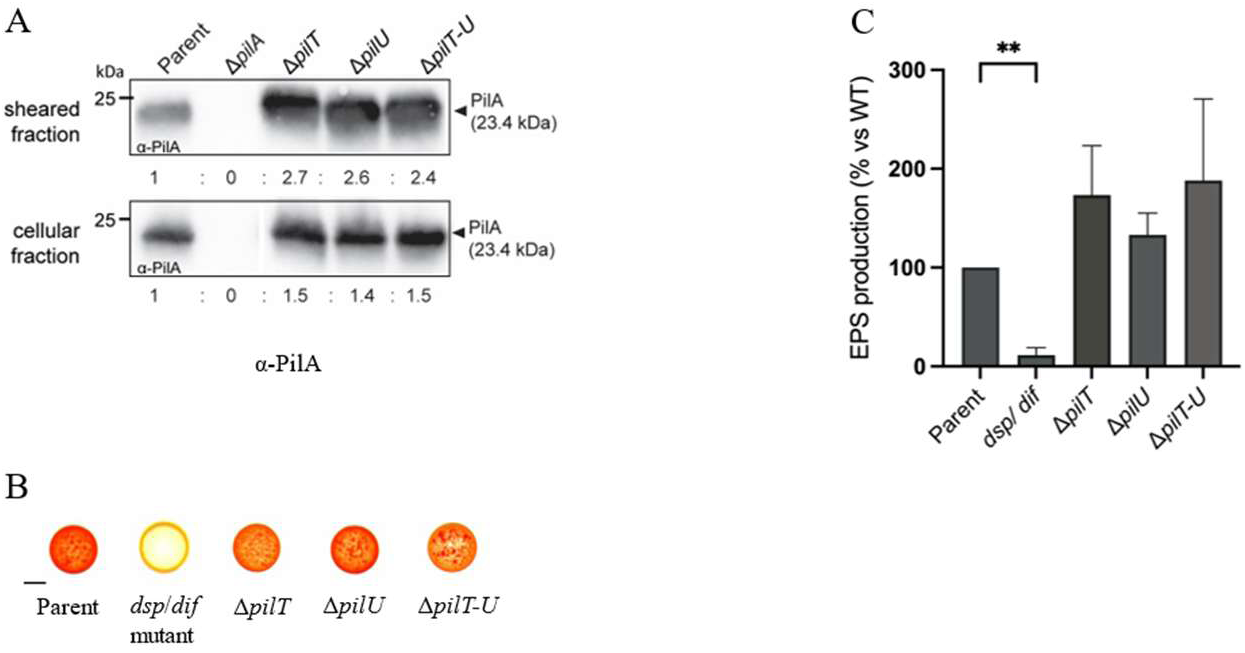
Pili and exopolysaccharide production in *pilT* and *pilU* mutants. (A) Shear-pili assay detecting PilA (24 kDa) by immunoblotting with polyclonal α-PilA antibody. Bands were quantified and relative to parent strain DK1217 were scored. (B) Congo red assay to assess exopolysaccharide (EPS) production; *dsp/dif* mutant serves as EPS-deficient control (yellow-orange color). Scale bar, 2.5 mm. (C) Quantitative trypan blue binding assay of EPS levels. Dye binding expressed as percentage relative to parent strain after subtraction of cell-free control. The double asterisk (**) denotes statistically significant differences in distributions (*p* < 0.0024) between parent strain and *dsp*/*dif* mutant.

Myxobacteria secrete EPS and fibril proteins, which together form an extracellular matrix used as an S-motility substrate. To check whether the PilU mutant makes EPS, we measured its production qualitatively and quantitatively with Congo red and trypan blue assays, respectively. After a 5-day incubation the *pilT*, *pilU* and *pilT-U* double mutants bound Congo red similar to the parent strain, as they produced bright red colonies. In contrast, a *dsp/dif* mutant [14, 35], defective in EPS production, served as negative control and produced yellow-orange colony due to low dye-binding capacity (Fig. 3B). To measure EPS levels, trypan blue binding assay was done where relative levels of EPS produced by the mutants were spectroscopically measured. The *pilU* mutant showed higher levels of EPS (120%) than the parent one, as did the *pilT* mutant (170%) and a double *pilT-U* mutant (180%) (Fig. 3C). Together, these data show that the Δ*pilU* mutant produces T4P and EPS, supporting the role of PilU in pili retraction.

### PilT and PilU form independent hexameric assemblies

To examine potential protein–protein interactions involving PilT and PilU, we employed the bacterial adenylate cyclase two-hybrid (BACTH) system, in which proteins of interest were co-expressed in an *E. coli cya* mutant as fusions to the T18 or T25 fragments of the *Bordetella pertussis* adenylate cyclase catalytic domain [36]. Interactions between fusion partners reconstitutes adenylate cyclase activity, leading to cAMP production and subsequent activation of the lactose or maltose operons. Protein–protein interactions were readily detected on indicator plates, where interacting fusion pairs produce blue-green colonies on X-gal plates, whereas non-interacting pairs yield colorless colonies (Fig. 4A).

**Figure 4.**
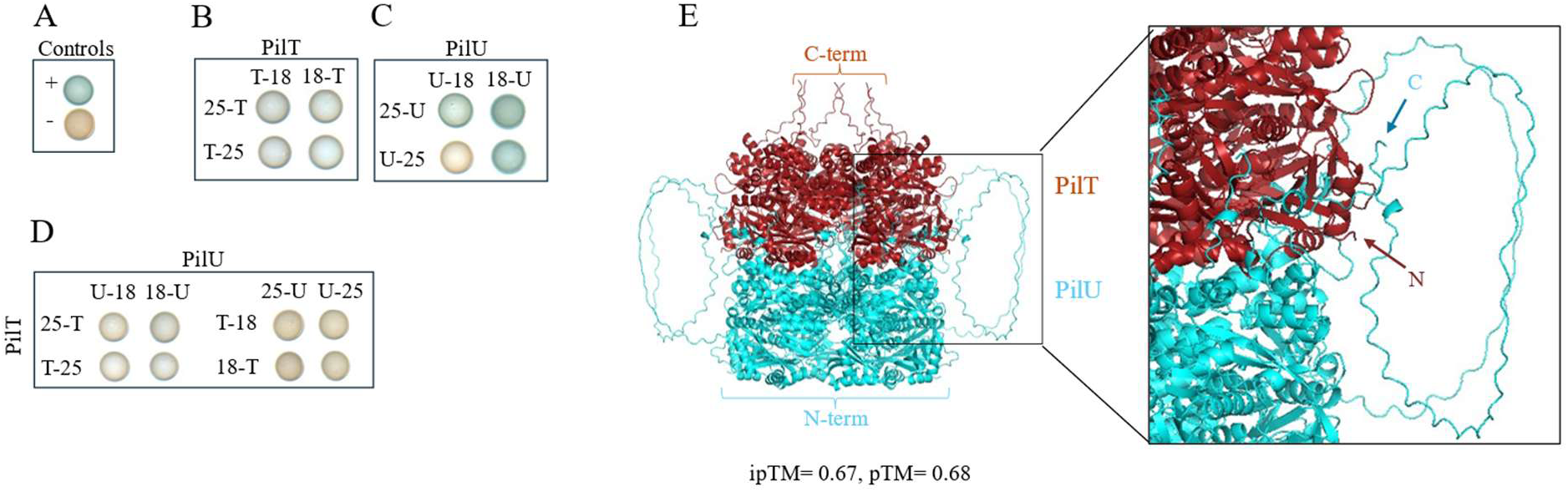
PilT and PilU do not interact as monomers but predicted to form co-structures of homohexamer rings. Bacterial adenylate cyclase two-hybrid (BACTH) assays testing pairwise interactions between T18- and T25-tagged PilT and PilU. (A) Leucine zipper fusions used as positive control and empty vectors as negative control. (B) PilT self-interaction. (C) PilU self-interaction. (D) Absence of detectable interactions between PilT and PilU. (E) AlphaFold3-predicted co-structure of PilT (red) and PilU (cyan). Visible N- and C-termini labeled.

Consistent with predicted hexameric architecture, BACTH analysis revealed strong self-interactions for both PilT and PilU (Fig. 4B, C). In contrast, no interaction was detected between PilT and PilU (Fig. 4D), suggesting that these fusion constructs do not form stable hetero-oligomeric complexes in the *E. coli* cytoplasm. Notably, AlphaFold predicts homo-oligomeric hexameric ring structures for PilT and PilU and, moreover, indicates a potential PilT–PilU association between their hexameric rings (Fig. 4E) where the N-terminal region of PilT binds to the C-terminal region of the PilU hexamer. The absence of detectable interaction in the BACTH assay suggests that assembled hexameric rings were a prerequisite for PilT and PilU associations that were likely transient.

### PilU localizes to the cell poles independently of PilT

Many key components of the T4PM, including the extension motor PilB and the major retraction motor PilT, localize to the cell poles, and several of these proteins undergo dynamic pole-to-pole oscillations during cellular reversals. More precisely, the PilB extension ATPase localizes to the leading cell pole, while PilT is asymmetrically bipolarly localized [19].

To determine whether PilU exhibits a similar localization pattern, we constructed a C-terminal GFP fusion using a monomeric superfolder GFP variant. The endogenous PilU-sfGFP fusion was functional (Fig. S4A), and it was stably expressed at the expected molecular mass (∼80 kDa) (Fig. S4B). Fluorescence microscopy revealed that PilU-sfGFP showed predominantly bipolar localization (63%) and unipolar localization (30%) (Fig. 5A, B). To directly compare the spatial distributions of the two retraction ATPases, we used a strain expressing an endogenous PilT-mCherry fusion. This endogenously expressed mCherry-PilT fusion was functional (Fig. S4A) and stably (∼87 kDa) (Fig. S4B). In comparison to PilU, PilT showed increased bipolar localization (∼90%) and decreased unipolar localization (10%) (Fig. 5A, B). We also constructed a strain expressing endogenously expressed PilU-sfGFP and mCherry-PilT and conducted colocalization studies. In this strain, both PilT and PilU showed unipolar and bipolar localization (Fig. 5A, B). Importantly, these localization distributions were similar whether PilT and PilU were expressed individually or simultaneously.

**Figure 5.**
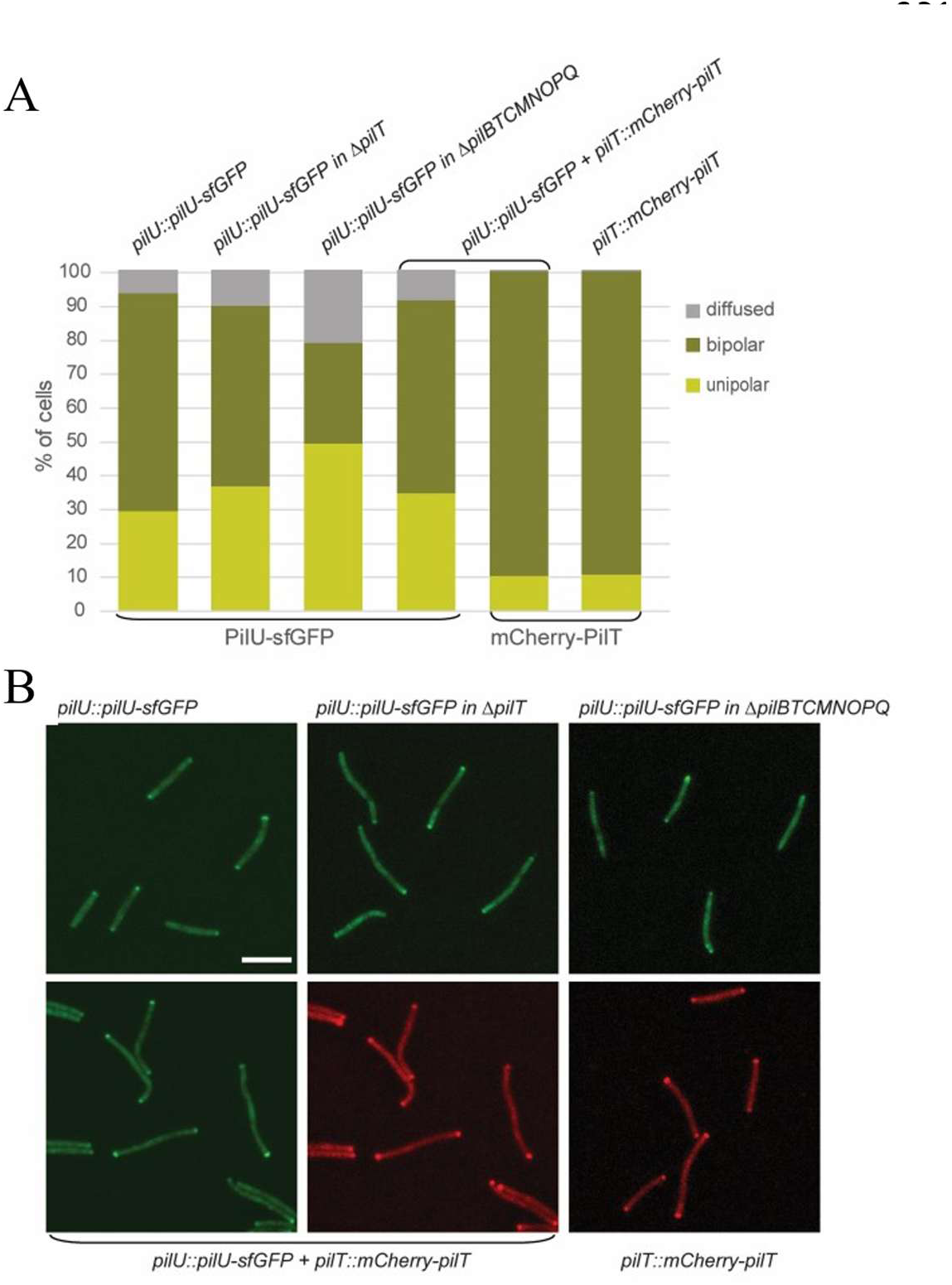
Polar localization of PilT and PilU. (A) Quantification of the polar localization of mCherry-PilT and PilU-sfGFP by fluorescence microscopy. (B) Micrographs show polar localization of PilU-sfGFP and mCherry-PilT. Scale bar, 5 µm.

Together, these results demonstrate that PilT and PilU were enriched at one or both cell poles, supporting the idea that the two retraction ATPases were spatially coordinated during T4P function. To determine whether PilU localization depends on PilT, we analyzed the localization of PilU-sfGFP in absence of *pilT*, which only resulted in minor changes to PilU-sfGFP localization (Fig. 5A, B), indicating that PilU localization was independent of PilT, suggesting that PilU was recruited to the cell poles by a distinct targeting mechanism. In contrast, expressing PilU-sfGFP in a strain lacking the core T4P assembly components (Δ*pilBTCMNOPQ*) produced a 3-fold increase in diffuse cytoplasmic localization, a ∼2-fold increase in unipolar localization and a concomitant reduction in bipolar polar localization (Fig. 5A, B). These shifts suggest that although PilT was not required for PilU polar targeting, components of the core pilus assembly machinery or an extended T4P machine contribute to the stable polar enrichment of PilU. Finally, we note that overexpression of *pilU* in a *pilT-U* double mutant failed to restore motility, further demonstrating the central role PilT plays in pili retraction and motility (Fig. S5).

### PilU contributes to efficient single-cell T4P–dependent motility

To assess the contribution of PilT and PilU to single-cell motility, we performed time-lapse microscopy in strains that lacked A-motility (Fig. 6, supplementary videos 1-4). For this, strains were grown to mid-log phase and spotted on 0.5% agar pad in 0.5% CTT supplemented with 2 mM CaCl_2_ and incubated at 33 °C for 1 hr in a humid chamber, then 100 individual cells were tracked over a 30 min period. As expected, the control cells (DK1217) displayed robust motility, with nearly all tracked cells exhibiting persistent movement during the image period. In contrast, the *pilT* mutant was completely nonmotile, consistent with the established role of PilT as the primary T4P retraction motor. The Δ*pilU* mutant resulted in a strong but incomplete motility defect. While the majority of Δ*pilU* cells remained immobile, a small subpopulation (∼10%) displayed detectable movement during the 30-min observation window. This residual motility was infrequent and characterized by short-lived movements with minimal net displacement, which supports no net migration of *pilU* mutants found on soft agar swarm plates.

**Figure 6.**
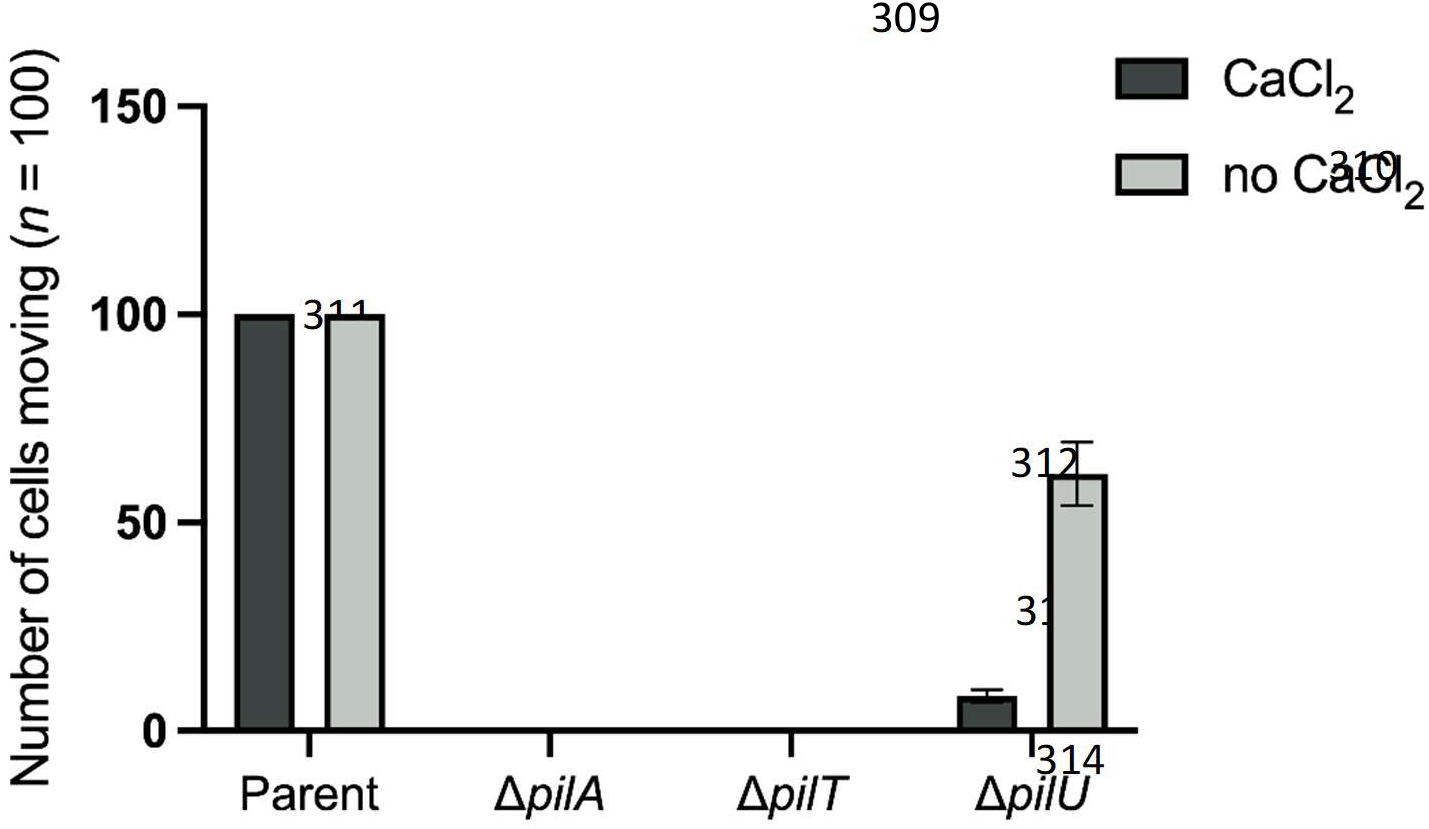
Calcium modulates *pilU*-independent motility. Single-cell motility of the parent strain and *pil* mutants in the presence and absence of 2 mM Ca²⁺. Time-lapse microscopy done over 30 min with images acquired every 16 s. For each strain, 100 individual cells were tracked. Bar graph shows the number of cells that move at least once over a 30 min duration.

### Calcium differentially modulates PilU-dependent motility

Calcium is a known modulator of S-motility [37–40], and to assess the influence of this cation, we examined colony swarming of WT, Δ*pilT*, Δ*pilU*, and Δ*pilA* strains on CTT soft agar in the presence and absence of CaCl₂ in both A^+^ and A^—^ backgrounds. In an A^+^ background in the absence of CaCl₂, the WT strain exhibited large irregular S-motility flares, while the Δ*pilU* mutant showed limited spreading accompanied by short, irregular flares at the colony edge (Fig. S6A). In the presence of CaCl₂ WT motility was enhanced in that colony expansion was evenly dispersed, while the Δ*pilU* colony showed reduced expansion and was also more evenly dispersed compared to the presence of CaCl₂ (Fig. S6A).

In A^—^ backgrounds in the absence of CaCl₂, the parent A^—^S^+^ strain exhibited irregular S-motility flares with moderate colony expansion, while in the presence of calcium it showed an enhanced and evenly dispersed colony morphology (Fig. S6B). In contrast, in the presence of calcium the swarming phenotype of the Δ*pilU* mutant was completely abolished (Fig. S6B). Given the opposite effects of CaCl₂ on the parent and Δ*pilU* phenotypes, we next examined calcium-dependent effects on motility at the single-cell level in an A^—^ background. For the parent strain, time-lapse microscopy revealed that CaCl₂ did not change the number of cells moving (Fig. 6, supplementary video 5; compare to supplementary video 1). In contrast, Δ*pilU* cells exhibited very limited movement in the presence of CaCl₂, with only a small fraction of cells showing short, infrequent displacements. However, in the absence of CaCl₂, motility of the Δ*pilU* mutant was substantially increased, with approximately ∼55% of cells displaying detectable movements that were more frequent and sustained than under calcium-supplemented conditions (Fig. 6, supplementary video 4; compare to supplementary video 8). For *pilA* and *pilT* mutants, there were no cell movements in the presence or absence of CaCl_2_ (Fig. 6, supplementary video 6, 7; compared to supplementary video 2, 3). Together, these results indicate that calcium differentially modulates T4P-driven motility of *pilU* mutants in the absence or presence of the A-motility gliding machinery, revealing an important role of PilU in supporting S-motility under calcium-rich conditions.

### Suppressor reveals a regulatory role for the PilU C-terminus

To gain PilU functional insights, we screened for spontaneous suppressors of the *pilU*-*fs* mutant that restored motility. A suppressor was isolated and whole-genome sequencing revealed an intragenic nonsense mutation near the 3′ end of the *pilU-fs* mutation. This mutation, Q405→NS, truncated PilU from its full length of 468 amino acids to 404 amino acids. Backcrossing this nonsense mutation into the original *pilU*-*fs* background restored motility, demonstrating that this mutation conferred suppression (Fig. 7A). Structural predictions with AlphaFold indicated that PilU contains a flexible, intrinsically disordered C-terminus (Fig. 7B). The 2120 *pilU-fs* mutation at codon 454 extended the C-terminus by 36 amino acids (Fig. 7C), whereas the suppressor mutation shortened the WT protein by 64 amino acids and essentially removed the intrinsically disordered C-terminus (Fig. 7D). These findings suggest that the frameshift mutation, which also elongates the C-terminus, interferes with PilU function, possibly by destabilizing hexamer formation, or by imposing a constitutive inhibitory effect. In the suppressor, truncations of this IDR may facilitate proper hexamer assembly, or it removes a C-terminal negative regulatory element, thereby restoring PilU function and motility.

**Figure 7.**
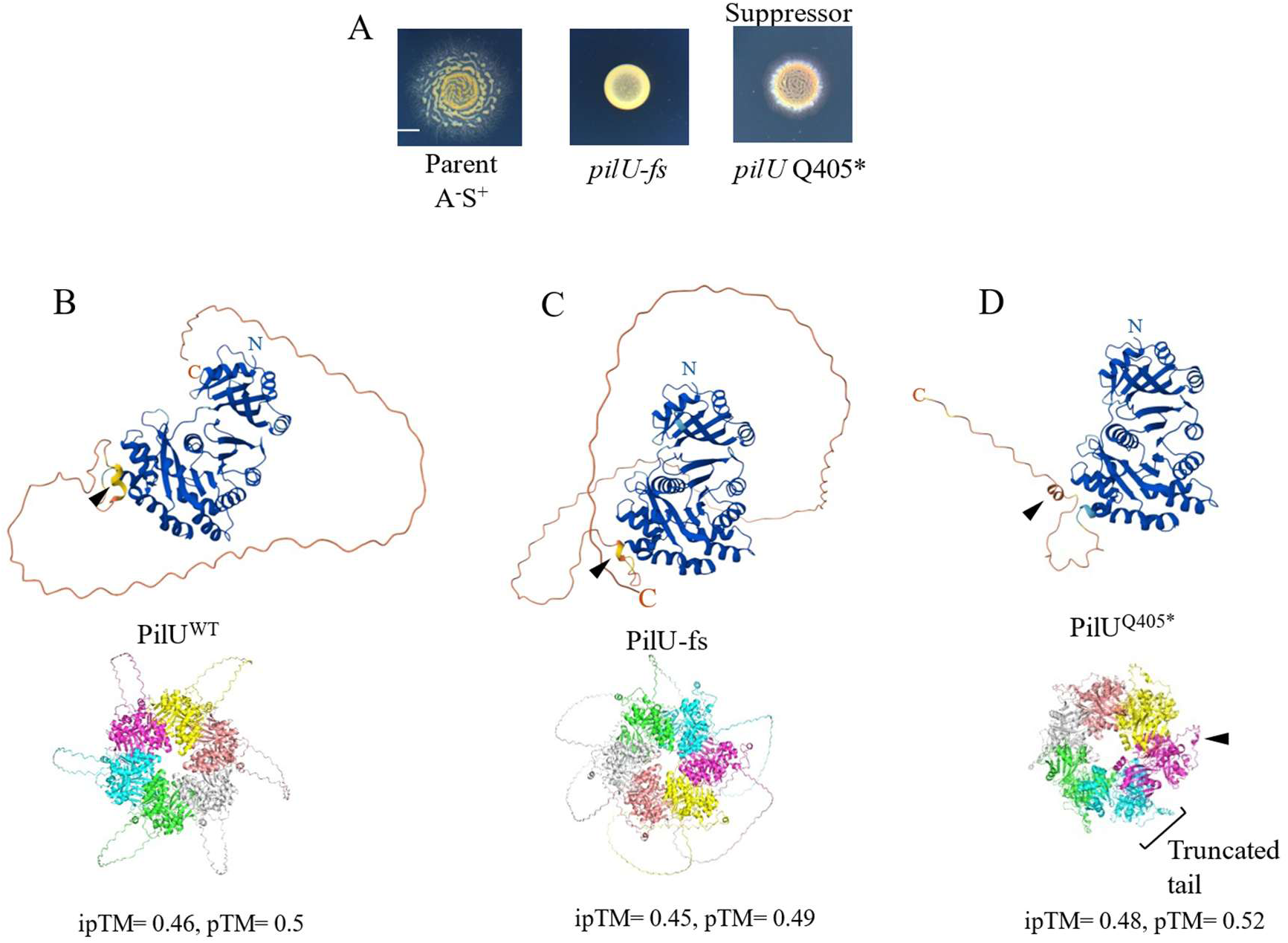
Intragenic suppression of the *pilU-fs* 2020 allele. (A) Colony phenotypes of the 2120 *pilU-fs* mutant and an intragenic suppressor carrying a nonsense mutation (Q405*). Scale bar, 2.5 mm. Parent strain DK1218. (B–D) AlphaFold models of the PilU monomer (upper) and hexamer (lower) for WT PilU (B), PilU-fs mutant with a 36 amino acid extended C-terminal tail (C), and the Q405* intragenic suppressor with a truncated C-terminal tail (D). For reference, black triangles mark the short helix near the C-terminus.

## Discussion

T4P–mediated motility in *M. xanthus* relies on tightly coordinated cycles of pilus extension, adhesion, and ATP-driven retraction. In this study, we identify PilU as an important yet previously unrecognized component of the T4P retraction machinery, along with PilT to power S-motility. Comparative genomics revealed that although *M. xanthus* encodes four PilT-like ATPases, deletion of *pilU* alone eliminates S-motility. Two of the remaining paralogs, *MXAN_0415* and *MXAN_6706*, exhibit partial S-motility defects that warrant further investigation. Despite PilU’s lower occurrence among myxobacteria species, orthologs are broadly found in other diverse T4P systems. Notably, *pilU* resides within a conserved gene cluster and, in related genera such as *Archangium* and *Melittangium*, it co-occurs with *pilY1* and minor pilin genes. These associated PilY1 proteins share sequence identity with the *M. xanthus* PilY1.3 variant [9], suggesting that *pilU* may have coevolved with specific minor pilin–PilY1 modules to fine-tune T4P-dependent motility.

Phenotypically, deletion of *pilU* closely resembles *pilT* loss-of-function at the colony level, producing smooth edged, nonexpanding colonies despite pilus production. However, unlike *pilT* mutants, which are immotile as single cells, *pilU* mutants retain limited surface motility. These movements are rare, transient, and result in negligible net displacement. In *M. xanthus,* PilT-independent retraction events occur at high velocity but low force [3], which likely explains the weak and nonproductive movements observed in *pilU* mutants, which is consistent with insufficient force generation despite rapid retraction dynamics. This residual activity also indicates that PilU is not the primary retraction motor but instead enhances or modulates PilT-dependent retraction, consistent with observations in *P. aeruginosa*, *V. cholerae*, and *Neisseria* spp [23–25], where PilU functions as an accessory retraction ATPase that depends on PilT and augments retraction force generation. Additionally, unlike *pilT* null mutants, Δ*pilU* mutants have conditional phenotypes where low temperatures or the absence of calcium partially restores motility when gliding is missing. Our prior studies with *sglS* and *sglT* null mutants also exhibit Ts^ꟷ^ S-motility phenotypes [31, 32], suggesting these two proteins may function with PilU in a similar pathway. Finally, the inability of PilU overexpression to restore motility in a *pilT pilU* double mutant further supports the central role PilT plays in motility.

Importantly, *pilU* deletion does not disrupt pilus assembly or EPS production, confirming that the S-motility defect is not the consequence of defective pilus assembly or matrix synthesis. Both *pilT* and *pilU* mutants displayed increased levels of surface-exposed pili and elevated EPS relative to the WT, suggesting the existence of feedback regulation between retraction efficiency and EPS secretion. This coordinated regulation likely serves to maintain optimal pilus–surface interactions required for effective T4P-mediated motility [41–44].

Our protein-protein interaction studies further clarify the relationship between PilT and PilU. Both ATPases exhibited self-interaction in the bacterial two-hybrid assay, consistent with the conserved hexameric architecture of T4P retraction motors. However, we did not detect a direct PilT–PilU interaction in a heterologous host, although our AlphaFold analysis predict these hexameric rings do interact. Experimental findings from other Gram-negative bacteria found that PilT and PilU interact or form mixed assemblies [23, 24, 45]. In *V. cholerae*, structural modeling, AlphaFold predictions, cryo-EM and molecular dynamics simulations have suggested that PilT and PilU homohexamers assemble into coordinated complexes and that interactions between PilT N-terminus and the PilU C-terminus are important for motor coordination during T4P retraction [46–48]. Our inability to detect such an interaction in BATCH assays suggests that coordination between PilT and PilU in *M. xanthus* may be transient, condition-dependent, or mediated by additional components of the pilus machinery.

Spatial localization of the motors provides additional insight into their functional relationship. In *M. xanthus*, the extension ATPase PilB localizes predominantly to the leading pole, whereas the retraction ATPase PilT is enriched asymmetrically at both poles [19]. When cells change their direction of movement, T4P assembles at the new leading pole, and together, PilB and PilT switch polarity. In contrast, studies in *P. aeruginosa* have shown that PilB and PilT can localize to both poles, while PilU is preferentially associated with the piliated pole [49]. In our study, PilU–GFP localized primarily to one or both poles, closely mirroring the distribution pattern of PilT. Notably, PilU remained polar, even in the absence of PilT, indicating that its localization does not depend on PilT. It is tempting to speculate that SofG and/or BacP are involved in the polar localization of PilU, as already demonstrated for PilB and PilT [50]. Frequent colocalization between PilT and PilU supports a model in which the two motors coordinate at the T4P base without forming stable hetero-hexamer complexes. The T4P machine at the leading pole can switch between an unpiliated state and an extension mode driven by PilB, while retraction is initiated when PilT replaces PilB to drive pilus depolymerization. We propose that, under high load, PilU acts with PilT to support high-force retraction, and in the absence of PilT, PilU may mediate low-force retraction (Fig. 8).

**Fig. 8.**
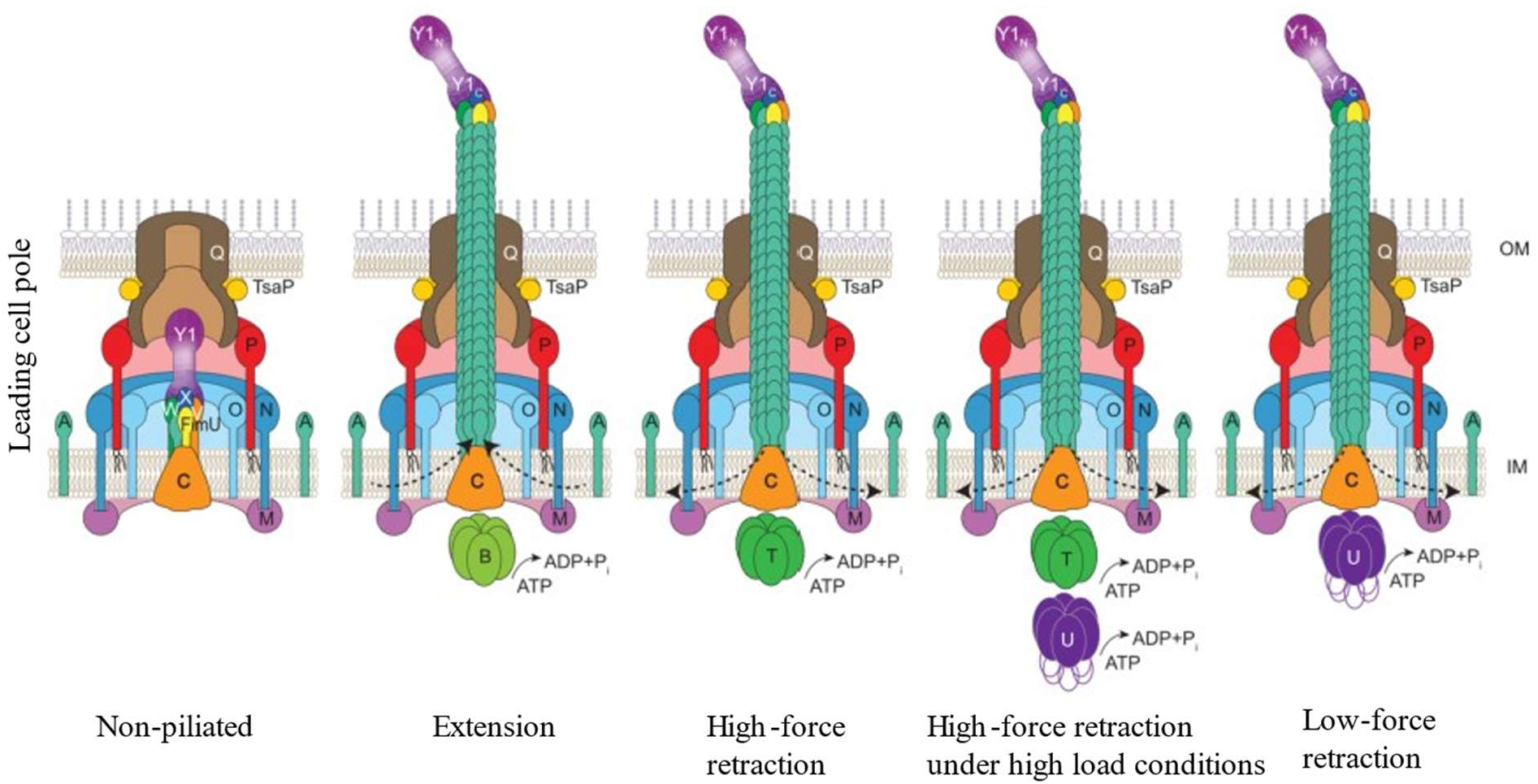
Model of T4PM at the leading cell pole in non-piliated and piliated states. (A) At the leading cell pole the T4PM can be unpiliated (left) or upon binding of PilB at the T4PM base the pilus polymerizes (2^nd^ left). For retractions, PilT replaces PilB and drives depolymerization of the pilus fiber into the cytoplasmic membrane (middle). Under high-load conditions PilU acts as an accessory retraction motor that together with PilT supports T4P retraction (2nd right). In absence of PilT high velocity but low-force retraction can occur, presumably by PilU. Dashed arrows indicate incorporation or removal of the major pilin PilA at the pilus base during extension and retraction, respectively. Single-letters denote canonical Pil-prefix designations of each protein. Y1_N_ and Y1_C_ denote the N- and C-terminal domains of PilY1.

A striking finding in this study is the identification of a spontaneous intragenic suppressor that restores motility to the *pilU* frameshift mutant. The suppressor introduces a nonsense mutation that truncates the C-terminal tail, including the 36 amino acid extension generated by the 2120 frameshift mutation. The PilU extended C-terminal tail is unique to myxobacteria, where AlphaFold modeling suggests it is intrinsically disordered, and the frameshift mutation further elongates this region, which may in turn disrupt hexamer stability or activate an auto-inhibition mechanism. Thus, the suppressor may relieve this interference, restoring functional hexamer formation, or alleviating a negative regulatory function. Consistent with the latter, IDRs are frequently involved in protein-protein interactions [51, 52], thus another factor may bind the IDR in PilU to regulate its function where SglS or SglT could play such a role. Regardless, these data underscore a role of the PilU C-terminus for proper motor activity, consistent with emerging evidence that flexible terminal regions of PilT-like ATPases modulate oligomer dynamics and ATPase cycling [15, 47, 48, 53].

Our data also reveal an unexpected calcium-dependent requirement for PilU. Whereas calcium enhances motility in WT cells, it abolishes the limited motility of Δ*pilU* mutants in the absence of A-motility. Calcium-dependent stabilization of adhesins and extracellular matrix components has been reported in other bacteria [54, 55], where it promotes surface attachment and early biofilm formation. Calcium binding also stabilizes PilY1 and ultimately allows minor pilin cluster to support T4P function in a broader range of environmental conditions [38, 39, 56]. Moreover, *M. xanthus* cells dynamically adjust the relative contributions of A- and S-motility in response to calcium, shifting from predominantly A-motility at low calcium to enhanced S-motility at higher calcium concentrations [40]. Our findings may reflect calcium-induced strengthening of pilus–EPS interactions that elevate mechanical load, rendering PilT alone insufficient for productive retraction. Alternatively, calcium may differentially modulate PilT and PilU activities or their integration with downstream signaling systems. Together, these observations suggest that extracellular calcium regulates the PilT–PilU retraction system, allowing environmental cues to fine-tune T4P-dependent motility by modulating retraction force and coordination.

Collectively, our results establish PilU as a functional PilT paralog and an accessory retraction motor that together with PilT supports high-force T4P retraction. This dual-motor system in *M. xanthus* reveals additional complexity in T4P regulation and suggests a mechanism for environmental control of retraction dynamics. Future studies integrating biophysics, structural analysis, and high-resolution imaging will help clarify how PilT and PilU coordinate force generation to drive multicellular behavior.

## Methods and Materials

### Bacterial strains and growth conditions

All bacterial strains used in this study are listed in Supplementary Table 1. *M. xanthus* cultures were grown in CTT medium (1% [w/v] casitone, 10 mM Tris–HCl pH 7.6, 1 mM KH₂PO₄, 8 mM MgSO₄) at 33 °C in the dark with shaking. *E. coli* strains were maintained in LB medium. When required, antibiotics were added at the following final concentrations: kanamycin (50 µg/mL), oxytetracycline (10 µg/mL), or ampicillin (100 µg/mL). Cells were washed in TPM buffer (CTT lacking casitone). Solid media were prepared by adding agar to 1.5% or 0.5%.

### Sequencing and genetic mapping

Genomic DNA from the DK2120 strain was sequenced using the Illumina NextSeq 2000 platform (MiGS, Pittsburgh, PA). Mutations were identified by aligning reads to the DK1622 reference genome using the BreSeq pipeline.

### Plasmid and strain construction

Plasmids and primers are listed in Supplementary Tables 2 and 3. For rescue experiments, the full-length *MXAN_1995* WT gene was amplified using Q5 high-fidelity DNA polymerase (NEB) and cloned into pMR3487 [57]. Constructs were propagated in *E. coli* DH5α, verified by colony PCR, restriction digestion, and sequencing, then electroporated into *M. xanthus* for chromosomal integration via homologous recombination under oxytetracycline selection.

Markerless deletions as well as integration of *pilU-sfGFP* and *mCherry-pilT* were generated using a two-step homologous recombination strategy [58]. Verified constructs were electroporated into *M. xanthus*, selected on kanamycin, then counter-selected for plasmid excision on 2% galactose–CTT agar [59]. Successful deletions and insertions were confirmed by PCR using primers flanking the deletion/insertion site and phenotypic analysis.

### Motility assays

*M. xanthus* cultures were grown in CTT to mid-log phase were washed with TPM buffer and normalized to 1.5 × 10⁹ cfu/mL. To assess S-motility, 5 µL of culture was spotted onto 0.5% agar CTT plates supplemented with 2 mM CaCl₂ (unless stated otherwise). Plates were incubated at 33 °C and imaged after 72 h using an Olympus SZX10 stereoscope.

For single-cell movement analysis, cells were spotted on 0.5% agar CTT glass slides containing 2 mM CaCl₂ (unless stated otherwise) and incubated at 33 °C for 1 h in a humid chamber. Time-lapse imaging was performed using an Olympus IX83 inverted microscope equipped with a 60× oil immersion objective and an Orca-Flash4.0 LT sCMOS camera.

### EPS assays

EPS production was examined by spotting 5 µL of cell suspensions (3 × 10⁸ cfu/mL in TPM) onto CTT plates containing 30 µg/mL Congo red. Plates were incubated at 33 °C for 5 days before imaging [32]. Retention of trypan blue was carried out as previously described [60]. Cells from overnight cultures were resuspended in TPM to OD₆₀₀ = 1.0. A 900 µL aliquot was mixed with 100 µL of trypan blue (100 mg/mL). Samples were incubated for 1 h at room temperature with gentle rocking, then centrifuged at 16,000 × g for 5 min. The top 900 µL of supernatant was transferred to cuvettes, and absorbance at 585 nm was measured. Values were normalized to WT levels.

### Fluorescence microscopy and image analysis

Fluorescence microscopy was performed as described earlier [19]. Briefly, exponentially growing cells were placed on a coverslip and overlaid with an agarose pad (1% SeaKem LE agarose (Cambrex) with TPM buffer (10 mM Tris-HCl pH 7.6, 1 mM KH_2_PO_4_ pH 7.6, 8 mM MgSO_4_) and 0.2% CTT). After 30 min at 32°C, cells were visualized using a temperature-controlled Leica DMi8 Thunder imager inverted microscope and phase contrast and fluorescence images acquired using a Leica K8 CMOS-camera and the LASX software (Leica Microsystems). Microscope images were processed with Fiji [61] and cell masks determined using Oufti [62] and manually corrected when necessary. To precisely quantify the localization of fluorescently labeled proteins, we used Matlab R2020a (The MathWorks) in an established analysis pipeline [63].

### Pili shear assay

Pili were sheared from *M. xanthus* cells using an established protocol [19]. Briefly, pili were isolated from *M. xanthus* cells grown on 1% CTT/1.5% agar for 2–3 days. All strains were the DK1217 parent background, except the negative control strain SA7705, which is Δ*pilA ΔaglQ*. Cells were scraped and resuspended in pili resuspension buffer (100 mM Tris-HCl pH 7.6, 150 mM NaCl; 1 mL per 60 mg cells). Suspensions were vortexed at maximum speed for 10 min. A 100 µL aliquot was collected for whole-cell lysates. Remaining suspension was centrifuged at 13,000 × g at 4 °C, and supernatants were clarified by two additional rounds of centrifugation. Pili were precipitated using 10× pili precipitation buffer (final concentrations: 100 mM MgCl_2_, 500 mM NaCl, and 2% PEG 6000) for overnight at 4 °C, then resuspended in SDS lysis buffer for SDS-PAGE.

### Immunoblotting

Whole-cell fractions or isolated pili were separated by SDS-PAGE. Rabbit primary antibodies included α-PilA, α-mCherry, and α-GFP. HRP-conjugated goat anti-rabbit secondary antibody (1:15,000) was used for detection. Blots were developed using HRP substrate (Millipore) and imaged with a BioRad ChemiDoc MP image analyzer.

### BACTH assays

BACTH assays were performed according to the manufacturer’s protocol (Euromedex) [36]. Briefly, plasmids encoding full-length PilT and PilU fused N-terminally or C-terminally to the T25 or T18 *B. pertussis* adenylate cyclase (CyaA) fragments were transformed into *E. coli* BTH101 alone or in pairs. As a positive control, BTH101 co-transformed with the plasmids pKT25-zip and pUT18C-zip. Transformed cells were incubated at 30 °C for 24 h, outgrown with 500 μL LB for 1 h at 37 °C with shaking. Transformants were isolated on LB plates containing kanamycin 50 μg/mL and ampicillin 100 μg/mL to select for both plasmids. Strains were then grown statically at 30 °C from frozen stocks overnight in LB supplemented with kanamycin and ampicillin and 3 μL of each culture was then spotted onto LB plates containing kanamycin, ampicillin, 0.5 mM IPTG and 40 μg/mL X-gal and incubated at 30 °C for 48 h.

### AlphaFold-Multimer modeling

The PilU structural model was generated using AlphaFold 3 [64]. PyMOL v2.4.1 (Schrödinger LLC) was used for visualization and analysis.

### Bioinformatics Analyses

Protein co-occurrence with PilU was examined using STRING [65]. Sequence alignments were produced with ClustalOmega [66]. Protein domains were annotated using InterPro. Phylogenetic trees were generated in MEGA7 using the Neighbor-Joining method.

## Acknowledgments

This work was supported by the National Institutes of Health grant GM140886 to D.W.

